# Early Epigenetic and Metabolic Responses to the Adipocyte Secretome Reveal Stress-Adaptive States in Triple-Negative Breast Cancer

**DOI:** 10.64898/2026.04.06.716548

**Authors:** Ashley Townsel, Maya Jaffe, Shiyu He, Yifei Wu, Asia Ingram, Madison Tipton, Melissa Kemp, Curtis J. Henry, Karmella A. Haynes

## Abstract

Obesity is a well-established risk factor for triple-negative breast cancer (TNBC), yet how adipocyte-derived signals reprogram cancer cell metabolism and chromatin states remains poorly defined. Here, we investigate how adipocyte-driven lipogenesis reshapes metabolic-epigenetic coupling to support stress-adaptive cell states and functional changes in epithelial TNBC cells. Using an integrated multi-omic approach, we combine RNA sequencing (RNA-seq), chromatin accessibility (ATAC-seq), metabolic flux modeling, and functional metabolic assays in lipogenic BT-549 cells. Computational modeling trained on RNA-seq predicts shifts in metabolic pathway usage, including enhanced NAD-linked metabolism. RNA-seq reveals a predominance of gene activation, consistent with ATAC-seq data showing a strong bias toward increased accessibility. Regions of increased accessibility are enriched for stress-adaptive and antioxidant pathways, including superoxide dismutase 2 (*SOD2*) and metallothioneins (*MT1F*, *MT1E*, *MT2A*). Functionally, lipogenic cells exhibit increased spare respiratory capacity, altered ATP-linked respiration, elevated extracellular acidification, and reduced reactive oxygen species (ROS) accumulation, consistent with a bioenergetically flexible, stress-adaptive metabolic state. Together, these findings reveal that adipocyte-driven metabolic rewiring promotes selective chromatin opening and activation of stress-adaptive gene programs, enabling TNBC cells to buffer oxidative pressure for enhanced proliferation and survival after exposure to the adipocyte secretome.

## INTRODUCTION

Adipocytes are abundant constituents of the breast tumor microenvironment and exert strong paracrine influence on epithelial cells through endocrine, inflammatory, and metabolic signaling. These effects appear to be subtype-specific: while adipose tissue promotes receptor-positive (ER+) tumor biology largely through endocrine estrogen signaling (1), accumulating evidence indicates that triple negative breast cancer (TNBC) cells mount a pronounced transcriptional and phenotypic response to adipocyte-derived inflammatory and metabolic cues (2–5). TNBC is disproportionately diagnosed in patients with obesity (body mass index >30 kg/m^2) (6,7) and exhibits impaired chemotherapy response in this population (8), suggesting that adipocyte signaling may reinforce stress tolerance and aggressive phenotypes in ER-negative disease. The TNBC subtype is disproportionately diagnosed among Black women, who have the lowest 5-year survival rate compared to other populations (9). Despite strong epidemiologic and preclinical support for a link between obesity and TNBC progression, the molecular principles by which adipocyte signals are integrated into sustained cancer cell states, particularly at the level of chromatin regulation, remain unresolved.

Experimental models have shown that adipocyte-rich microenvironments promote TNBC proliferation, invasion, and therapy resistance through exposure to cytokines, lipids, and extracellular vesicles (10–12). At the cellular level, adipocyte signaling drives extensive metabolic remodeling, including increased lipid uptake and synthesis, altered glycolytic flux, and enhanced mitochondrial and peroxisomal oxidative pathways (13–16). These changes support survival under metabolic and chemotherapeutic stress (15,17,18) but simultaneously impose risks associated with lipotoxicity and reactive oxygen species (ROS) accumulation. Lipogenic cancer cells mitigate these risks by reprogramming lipid composition, sequestering polyunsaturated fatty acids in lipid droplets, and engaging antioxidant pathways that buffer oxidative damage (19,20). Such adaptations enable cancer cells to tolerate otherwise deleterious metabolic states and are increasingly recognized as contributors to drug resistance (21). However, while these metabolic phenotypes are well documented, how adipocyte-induced metabolic rewiring is stabilized or maintained at the level of gene regulation remains poorly defined.

Metabolic rewiring has the potential to influence chromatin regulation by altering the availability of metabolites that serve as cofactors for epigenetic enzymes. For instance, histone lysine acetylation, deacetylation, methylation, and demethylation rely on metabolite pools such as acetyl-CoA, S-adenosylmethionine, α-ketoglutarate, and FAD, which are directly linked to lipid metabolism and mitochondrial function. Dysregulation of chromatin modifiers, including histone deacetylases, histone methyltransferases, and Polycomb-associated enzymes, has been widely implicated in TNBC progression (22,23). Obesity-associated signaling has been reported to both repress and activate chromatin through changes in enzyme activity and metabolite availability, suggesting that metabolic state can bias epigenetic remodeling (24). Metabolic modeling approaches, including those applied in this study, predict that lipogenesis and oxidative rewiring constrain the availability of epigenetic cofactors, raising the possibility that metabolic state imposes limits on chromatin remodeling capacity (25–28). How such metabolic constraints shape genome-wide chromatin accessibility and transcriptional responses, and how these changes impact the function of TNBC cells exposed to the adipocyte secretome is an area of active investigation.

Here, we integrate functional assays, systems-level metabolic modeling, transcriptomics, and chromatin accessibility profiling to investigate how adipocyte-mediated signaling reshapes metabolic-epigenetic coupling in TNBC cells. We find that transient exposure to adipocyte-conditioned medium induces a sustained growth phenotype, consistent with the establishment of durable cellular states rather than transient signaling effects. Computational modeling trained on RNA-sequencing data reveals adipocyte-driven metabolic rewiring characterized by enhanced NAD-linked metabolism and pathways associated with oxidative stress tolerance. Chromatin accessibility profiling at an early time point (24 hours) demonstrates that these metabolic changes are accompanied by asymmetric chromatin remodeling, with a strong bias toward increased accessibility across gene-proximal regulatory regions. Genes associated with increased accessibility are enriched for pathways involved in stress balance and antioxidant defense, consistent with selective activation of adaptive programs. Notably, comparatively limited decreases in accessibility at this stage suggest that transcriptional repression may arise through mechanisms that do not require widespread chromatin closure or may emerge over longer timescales. Functional metabolic analyses further show that adipocyte-exposed cells exhibit increased spare respiratory capacity, altered ATP-linked respiration, and reduced ROS accumulation, consistent with a bioenergetically flexible, stress-adaptive metabolic state. Together, these findings reveal that early adipocyte-driven metabolic rewiring promotes selective chromatin opening and activation of stress-adaptive gene programs, while pointing to distinct regulatory pathways governing gene repression, enabling TNBC cells to buffer oxidative pressure and sustain proliferation.

## RESULTS

### Transient Exposure to Adipocyte Conditioned Medium Induces a Sustained Growth Phenotype in TNBC Cells

To determine whether adipocyte-derived signals exert lasting effects on TNBC proliferative potential, we performed clonogenic colony formation assays following either constant or transient exposure to adipocyte-conditioned medium (ACM). Human BT-549 and HCC1806 TNBC cell lines, as well as the murine TNBC cell line 4T1, were cultured in unconditioned medium (UCM) or ACM for three days, harvested, and reseeded at low density to assess long-term colony growth over a three-week period (**Fig. 1A**). Across all three cell lines, continuous exposure to ACM significantly increased colony size compared to UCM controls, consistent with a pro-growth effect of adipocyte-derived signals (**Fig. 1B**).

**Figure 1.**
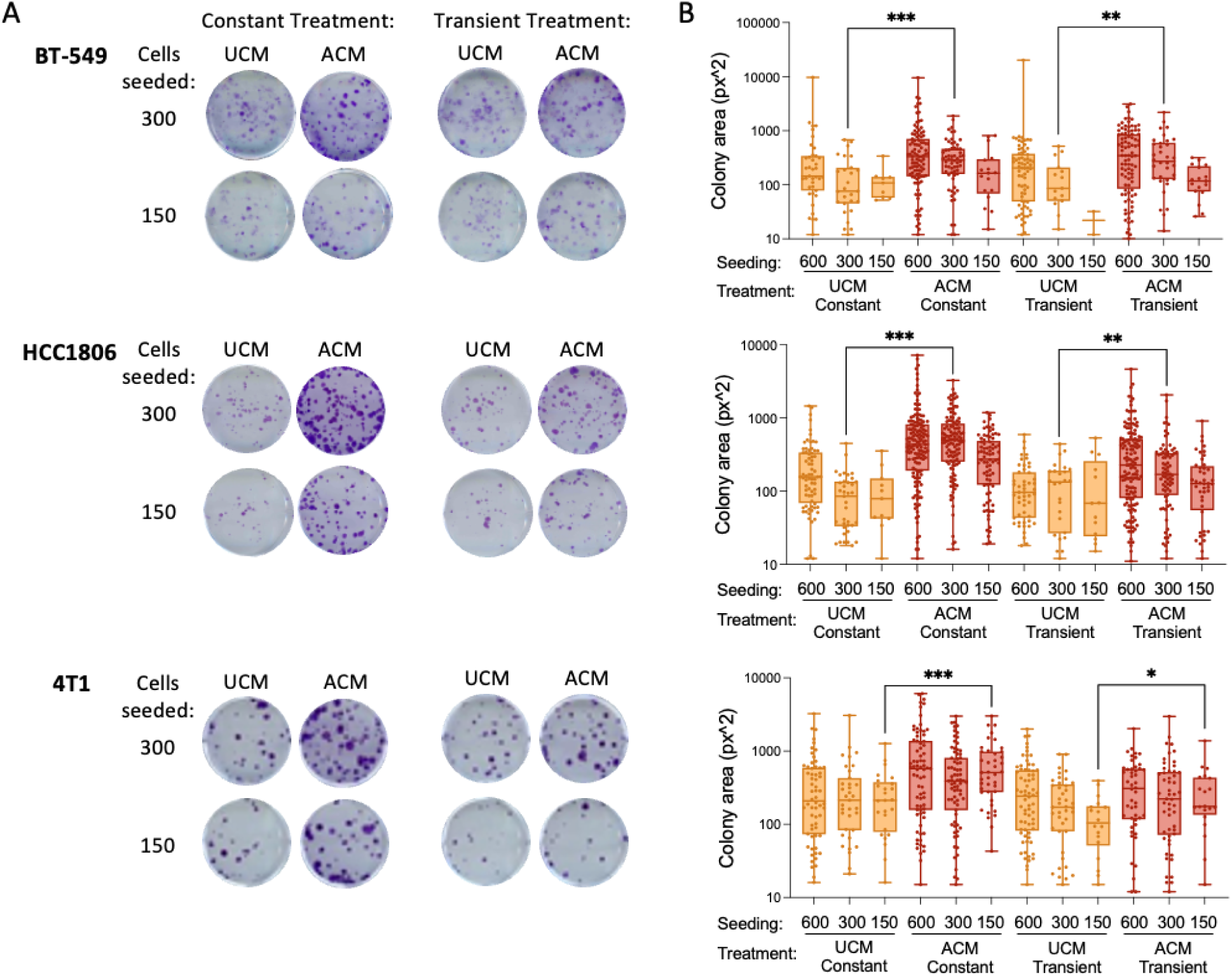
Changes in proliferation determined by colony formation assays for three TNBC cell lines. (A) Crystal violet staining of colonies grown from BT-549, HCC1806, or mouse 4T1 TNBC cells treated with unconditioned medium (UCM) or 100% adipocyte conditioned medium (ACM). Cells were grown with UCM or ACM for 3 days, harvested, then seeded at 300 or 150 per well and allowed to form colonies for 3 weeks. For the ACM-treated cells “constant” indicates colony formation in ACM for the duration of the experiment, and “transient” indicates colony formation after removal of ACM and the addition of UCM. (B) Box plots show quantification of clonogenic assay images (1 well per condition). *p* < 0.05*, 0.01**, 0.001*** (Student’s t-test, unpaired, two-tailed).

Notably, cells transiently exposed to ACM prior to reseeding also exhibited significantly enhanced colony growth relative to UCM-treated cells, despite removal of ACM during the colony formation phase. This sustained increase in clonogenic growth was observed at both seeding densities tested (300 and 150 cells per well), indicating that the effect is robust and not restricted to a narrow plating range. While the magnitude of the response varied between cell lines, the qualitative pattern was conserved: transient ACM exposure was sufficient to induce a persistent growth advantage that was maintained over multiple weeks in standard growth conditions. These findings indicate that short-term exposure to the adipocyte secretome imprints a durable proliferative phenotype in TNBC cells, motivating subsequent analyses of chromatin, transcriptional, and metabolic states that could underlie this sustained response and impact TNBC cell proliferation and survival in a pro-inflammatory microenvironment.

### Systems Modeling Reveals Adipocyte-Driven Metabolic and Epigenetic Rewiring in Lipogenic TNBC Cells

Given the sustained augmentation on the proliferative capacity of TNBC cells after a brief exposure and removal of the adipocyte secretome, we hypothesized that adipocyte secreted factors extensively remodel the metabolic and epigenetic state of TNBC cells. Our model of obesity-associated TNBC involves the generation of lipogenic TNBC cells with ACM. When BT-549 triple negative breast cancer cells are stimulated with adipocyte secreted factors, they show increases in lipid droplet formation that mimic the phenotype of adipocytes (**Fig. 2A**). In our previous study (29), RNA-sequencing analysis comparing ACM- and UCM-treated cells after 24 hours of treatment revealed 552 upregulated and 244 downregulated genes (|fold change| ≥ 1.5; |log_2_FC| ≥ 0.58; *p* ≤ 0.05) (**Fig. 2B**). The differentially upregulated genes (UpDEGs) were enriched for pathways related to chemical responses, cell proliferation, cell migration, and lipid storage, whereas the differentially downregulated genes (DownDEGs) were primarily involved in cholesterol and alcohol metabolic processes.

**Figure 2.**
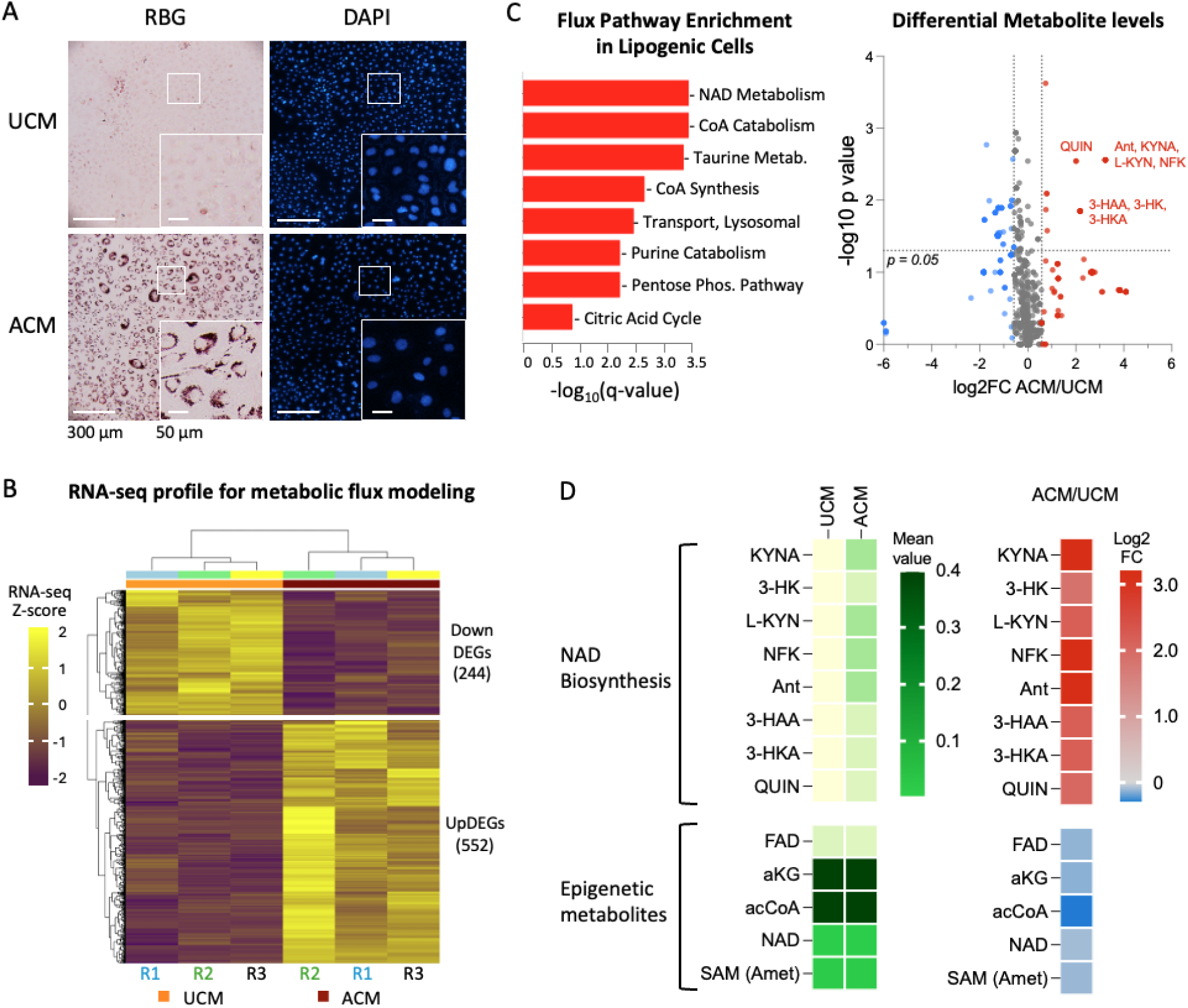
Adipocyte-conditioned media induces lipid accumulation and metabolic rewiring in human BT-549 cells. (A) Representative brightfield/RBG and DAPI images of BT-549 cells cultured in unconditioned media (UCM) or adipocyte-conditioned media (ACM), showing increased lipid droplet accumulation in ACM-treated cells. Insets show representative cells at higher magnification. Scale bars, 300 µm (main images) and 50 µm (insets). (B) Heatmap of z-scored RNA-seq expression values for differentially expressed genes (DEGs) identified between UCM- and ACM-treated cells (|log_2_FC| ≥ 0.58, *p* ≤ 0.05), used as input for metabolic modeling. (C) Differential metabolite predictions and pathway enrichment from genome-scale metabolic modeling constrained by RNA-seq data using the Recon3D human metabolic network. Left, pathway enrichment analysis of differentially altered flux-associated metabolites. Right, volcano plot of predicted metabolite changes in ACM relative to UCM, highlighting metabolites in the kynurenine-NAD biosynthesis axis. (D) Heatmap of selected modeled metabolites. Underlying values for significantly changed genes in panel C, and genes in panel D are provided in **Supplemental Table S1**.

To gain insights into metabolic and epigenetic changes at the onset of the lipogenic state in TNBC cells (24 hours after ACM treatment), we used the normalized gene expression values of our RNA-seq data to train a computational metabolic model. This human genome-scale metabolic model extends conventional Flux Balance Analysis (FBA) (30) by integrating quantitative, patient-specific, and cell-type-specific constraints, with a particular emphasis on redox metabolism relevant to cancer therapy. The framework is based on the Recon3D human metabolic reconstruction (31), comprising 8,401 metabolites and 13,547 reactions, which serves as the core metabolic network. Computational modeling predicted significant enrichment of NAD metabolism, along with CoA metabolism, taurine and hypotaurine metabolism, purine catabolism, the pentose phosphate pathway, lysosomal transport, and, to a lesser extent, the citric acid cycle (**Fig. 2C, Supplemental Table S1**). Inspection of differentially altered metabolites revealed a prominent increase in intermediates of the kynurenine pathway, including N-formyl-L-kynurenine (NFK), kynurenic acid (KYNA), 3-hydroxy-L-kynurenine (3-HK), 3-hydroxyanthranilate (3-HAA), and quinolinic acid (QUIN), consistent with activation of *de novo* NAD biosynthesis (**Fig. 2D**). Increased NAD biosynthesis is consistent with enhanced mitochondrial activity (32–35), which may support increased metabolic demands associated with proliferation in adipocyte-exposed TNBC cells (**Fig. 1**).

Metabolic pathways associated with redox balance were also prominently represented in the model. Elevated levels of kynurenine pathway intermediates and prostaglandins (**Supplemental Table S1**), which are associated with inflammatory processes and response to reactive oxygen species (ROS) (36–38), are consistent with a metabolic state that may generate oxidative pressure (39,40). In parallel, the model predicted increases in metabolites with reported antioxidant or cytoprotective functions, including xanthurenic acid (log2FC = 2.19, *p* = 0.01), which can limit lipid peroxidation (41,42), and sphingosine-1-phosphate (S1P, log2FC = 2.64, *p* > 0.05) (43,44). Together, these predictions suggest that adipocyte-exposed TNBC cells engage coordinated metabolic programs that balance redox-generating and redox-buffering processes, consistent with adaptation to a potentially oxidative microenvironment.

In contrast, metabolites commonly associated with chromatin regulation exhibited only modest changes in the model (**Fig. 2D**). Slight decreases were observed across cofactors including acetyl-CoA, α-ketoglutarate, NAD, SAM, and FAD, but these shifts were limited in magnitude and not statistically significant. These findings suggest that early lipogenic reprogramming does not substantially alter the steady-state availability of key epigenetic metabolites, but instead reflects regulatory mechanisms such as changes in metabolite flux, enzyme activity, or chromatin responsiveness that are not captured by bulk metabolite abundance. Consistent with this, metabolic changes were strongly centered on NAD-linked pathways and redox-associated processes, while canonical epigenetic cofactors exhibited comparatively modest shifts. Together, these results support a model in which chromatin regulation is shaped by localized or enzyme-specific dynamics rather than global changes in cofactor availability. To test whether these changes are reflected at the level of chromatin structure, we performed genome-wide chromatin accessibility profiling using ATAC-seq.

### Asymmetric Chromatin Remodeling Supports Selective Gene Activation in Lipogenic TNBC Cells

To investigate the impact of adipocyte-induced metabolic rewiring on chromatin accessibility, we performed ATAC-sequencing (ATAC-seq) on BT-549 cells treated with adipocyte-conditioned medium (ACM) or control medium (UCM). Differentially accessible regions (DARs) were identified using DiffBind (p value < 0.05, FDR < 0.05), revealing 3,764 regions with increased accessibility (UpDARs) and 710 regions with decreased accessibility (DownDARs) in ACM-treated cells. This distribution indicates that chromatin accessibility changes associated with the lipogenic state are predominantly skewed toward increased accessibility. Unsupervised clustering of the most significant DARs clearly separated ACM and UCM samples across biological replicates, demonstrating reproducible remodeling of the chromatin landscape (**Fig. 3A**). To further characterize these changes, we examined the distribution of log2 fold-change (log2FC) values across all differential peaks. The majority of DARs exhibited positive log2FC values, with relatively fewer peaks showing decreased accessibility (**Fig. 3B**). Accessibility changes were broadly biased toward modest increases (log2FC = 0.1 - 0.3) rather than large bidirectional shifts. We next annotated DARs by genomic location using ChIPseeker. Both UpDARs and DownDARs were enriched in genic regions, including promoters, introns, and exons, with comparatively fewer peaks located in distal intergenic regions (**Fig. 3C**). Intronic regions comprised the largest fraction of DARs, consistent with remodeling at gene-associated regulatory elements. These results indicate that the transition to a lipogenic state after 24 hours *in vitro* is accompanied by reproducible and predominantly positive shifts in chromatin accessibility, distributed across gene-proximal regions and characterized largely by modest effect sizes.

**Figure 3.**
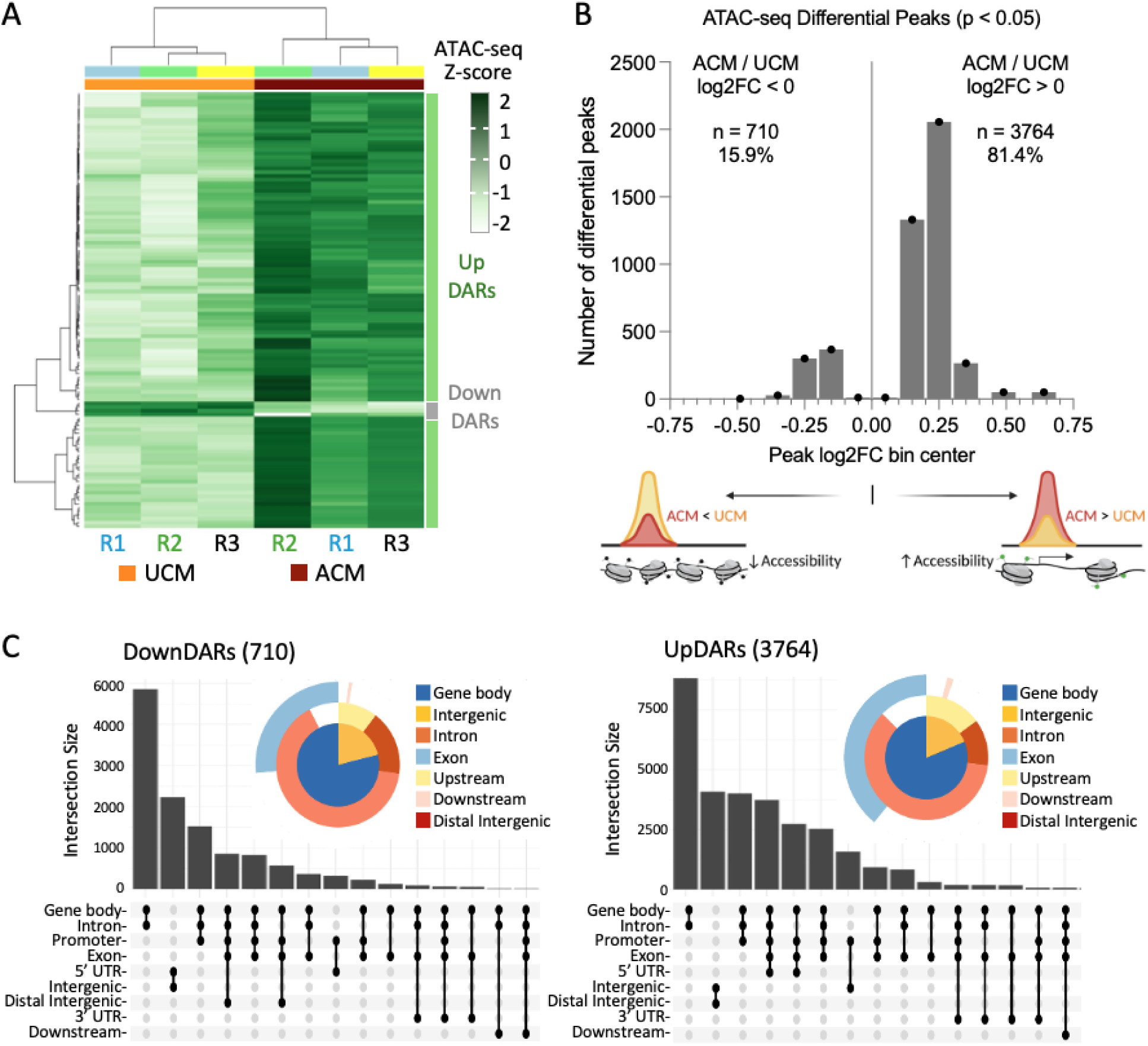
Global patterns of chromatin accessibility changes in lipogenic BT-549 cells. (A) Heatmap of the top 120 differentially accessible regions (DARs) identified by DiffBind (FDR < 0.05), ranked by adjusted *p*-value. Rows represent DARs and columns represent biological replicates (R1 - R3) from UCM- and ACM-treated cells. (B) Distribution of log2 fold-change (log2FC) values for ATAC-seq differential peaks (*p* < 0.05). Peaks are binned by log2FC and plotted as counts per bin. The majority of differential peaks exhibit positive log2FC values (UpDARs; *n* = 3764, 81.4%), whereas a smaller fraction show decreased accessibility (DownDARs; *n* = 710, 15.9%). (C) Genomic annotation of UpDARs and DownDARs using ChIPseeker. Donut plots summarize distribution across annotated genomic features. UpSet plots show intersections among annotation categories.

As a representative example of loci associated with repression, we examined the *PARVA* gene region, which harbors the insertion site of a transgenic reporter that becomes epigenetically silenced in lipogenic BT-549 cells (29). The *PARVA* locus exhibited relatively low baseline accessibility across intronic regions in both treatment conditions, with modest additional reduction in ACM-treated cells (**Supplemental Fig. S1**). Notably, the transgene insertion site resides within a chromatin environment that is comparatively inaccessible relative to an upstream region (*DKK3*, *MICAL2*). These observations suggest that, at least at this locus, transcriptional repression occurs within a chromatin context that is already relatively constrained rather than through dramatic *de novo* chromatin closure.

We next examined global histone acetylation. Immunoblot analysis revealed modest increases in H3K9ac and H3K27ac levels in ACM-treated cells relative to controls (**Fig. 4A**). These findings indicate that bulk histone acetylation is slightly elevated, suggesting the existence of an acetyl-CoA production mechanism. We therefore examined acetyl-CoA synthetase 2 (ACSS2), an enzyme that converts acetate to acetyl-CoA and has been reported to relocalize under metabolic stress (45). Although RNA-seq showed a modest, non-significant decrease in ACSS2 transcript abundance (log2FC = -0.45, *p* = 0.096), immunoblot analysis detected a predominant phospho-ACSS2 (pACSS2) species migrating at ∼100-150 kD in whole-cell lysates and in both cytoplasmic and nuclear fractions (**Fig. 4A**). Immunofluorescence microscopy revealed pronounced perinuclear enrichment of pACSS2 in ACM-treated cells compared to UCM controls (**Fig. 4B**). Quantitative single-cell fluorescence profiling demonstrated an approximately two-fold increase in the perinuclear-to-peripheral signal ratio (PNPR), consistent with redistribution of pACSS2 toward peri-nuclear regions under lipogenic conditions (**Supplemental Fig. S2**). This pattern may reflect association with endoplasmic reticulum-associated lipid droplets (46,47) and/or migration towards nuclear chromatin domains, consistent with a model in which localized acetyl-CoA production supports histone acetylation despite a lack of globally elevated acetyl-CoA predicted by our model (**Fig. 2D**).

**Figure 4.**
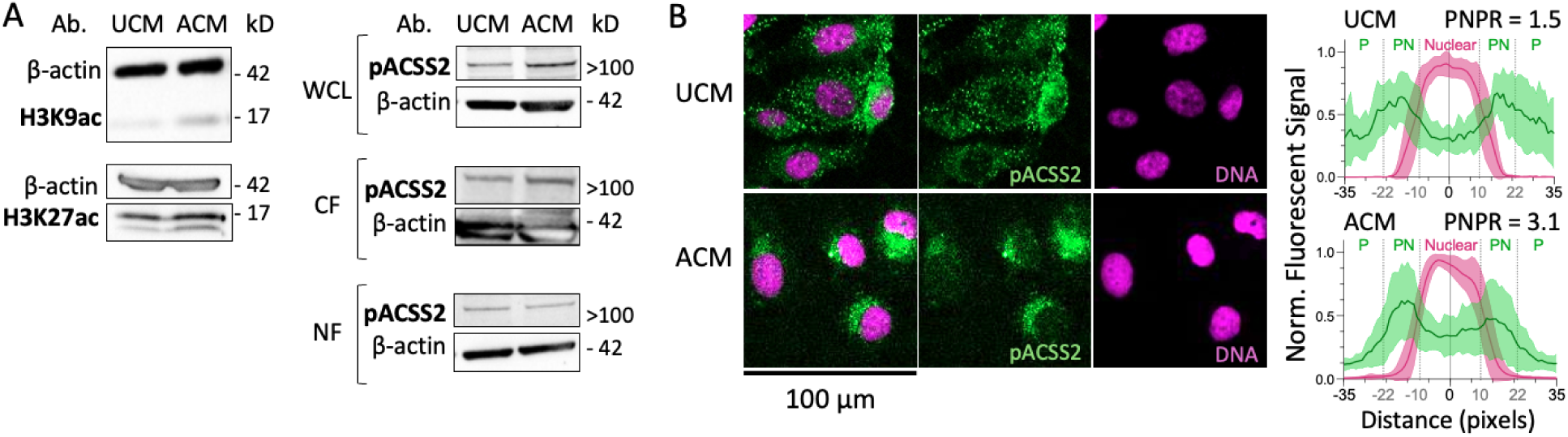
Maintenance of histone acetylation and redistribution of pACSS2 in lipogenic BT-549 cells. (A) Western blot analysis of H3K9ac, H3K27ac, and phospho-ACSS2 (Ser659) in control (UCM) and ACM-treated BT-549 cells. WCL, whole cell lysate; CF, cytoplasmic fraction; NF, nuclear fraction. (B) Left, immunofluorescence staining of UCM- and ACM-treated BT-549 cells for phospho-ACSS2 (Ser659) (green), with nuclear counterstaining using SiR-DNA (magenta). Right, single-cell radial fluorescence intensity profiles for each channel. Plots show mean signal (line) with standard deviation (shaded area), along with quantification of the perinuclear-to-peripheral ratio (PNPR) of pACSS2 signal (see **Supplemental Fig. S2**). *n* = 20 cells per condition.

To define the relationship between chromatin accessibility and transcriptional output, we integrated ATAC-seq and RNA-seq datasets by linking differentially accessible regions (DARs) to their associated genes. Upregulated genes (UpDEGs) were predominantly associated with regions of increased accessibility (UpDARs), with enrichment at promoter-proximal regions and gene bodies, consistent with a role for chromatin opening in facilitating transcriptional activation (**Fig. 5A**). Notably, increased accessibility was also observed at promoters and gene bodies of genes whose expression remained unchanged, suggesting that chromatin remodeling may precede or poise transcriptional responses in a subset of loci. In contrast, downregulated genes (DownDEGs) showed comparatively limited association with decreased accessibility (DownDARs), which were less frequent overall and more commonly localized to gene bodies rather than promoter regions. These patterns indicate that transcriptional repression in the lipogenic state is not broadly driven by promoter closure, but may instead reflect regulatory mechanisms that operate independently of large-scale accessibility changes.

**Figure 5.**
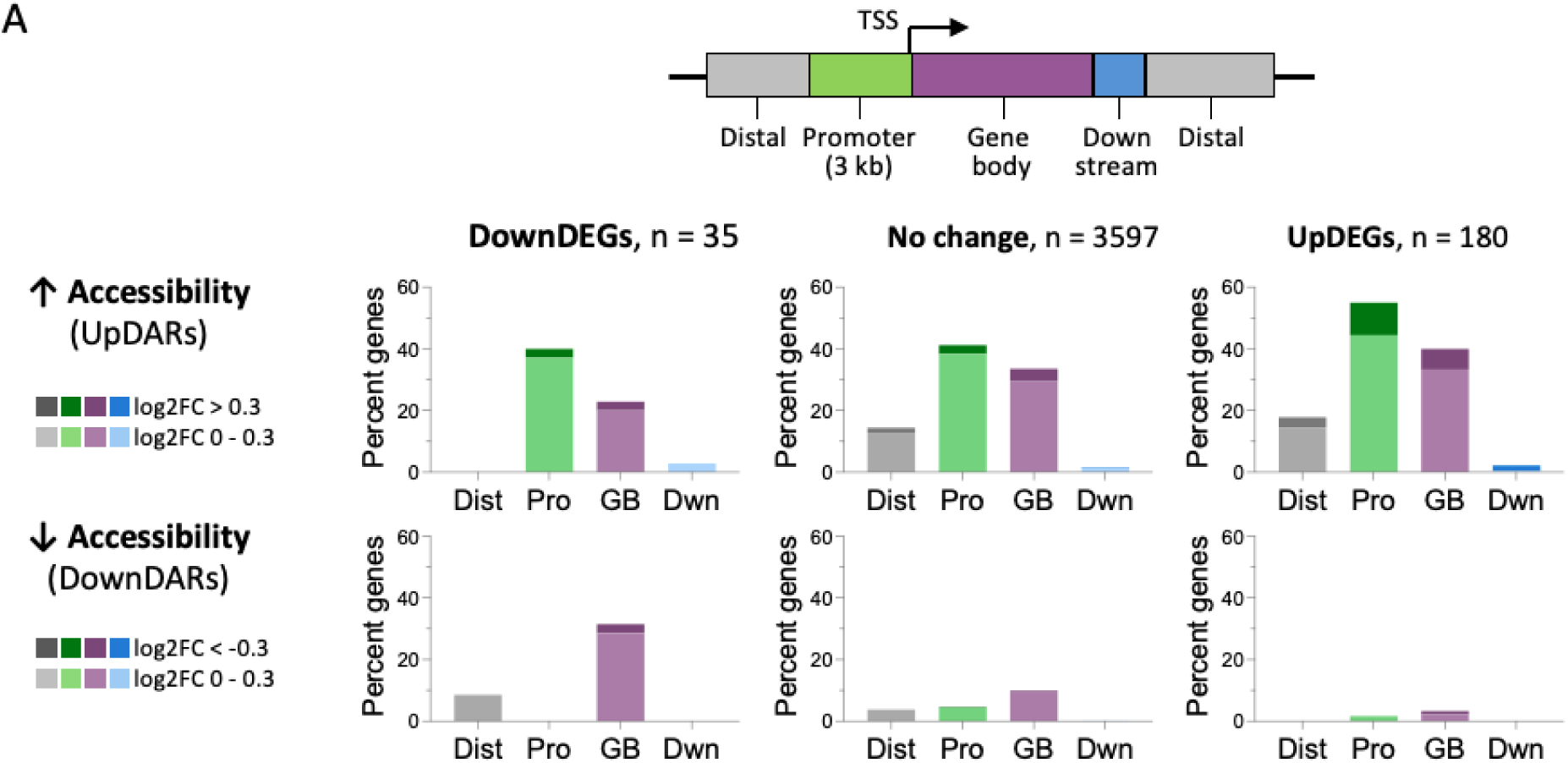
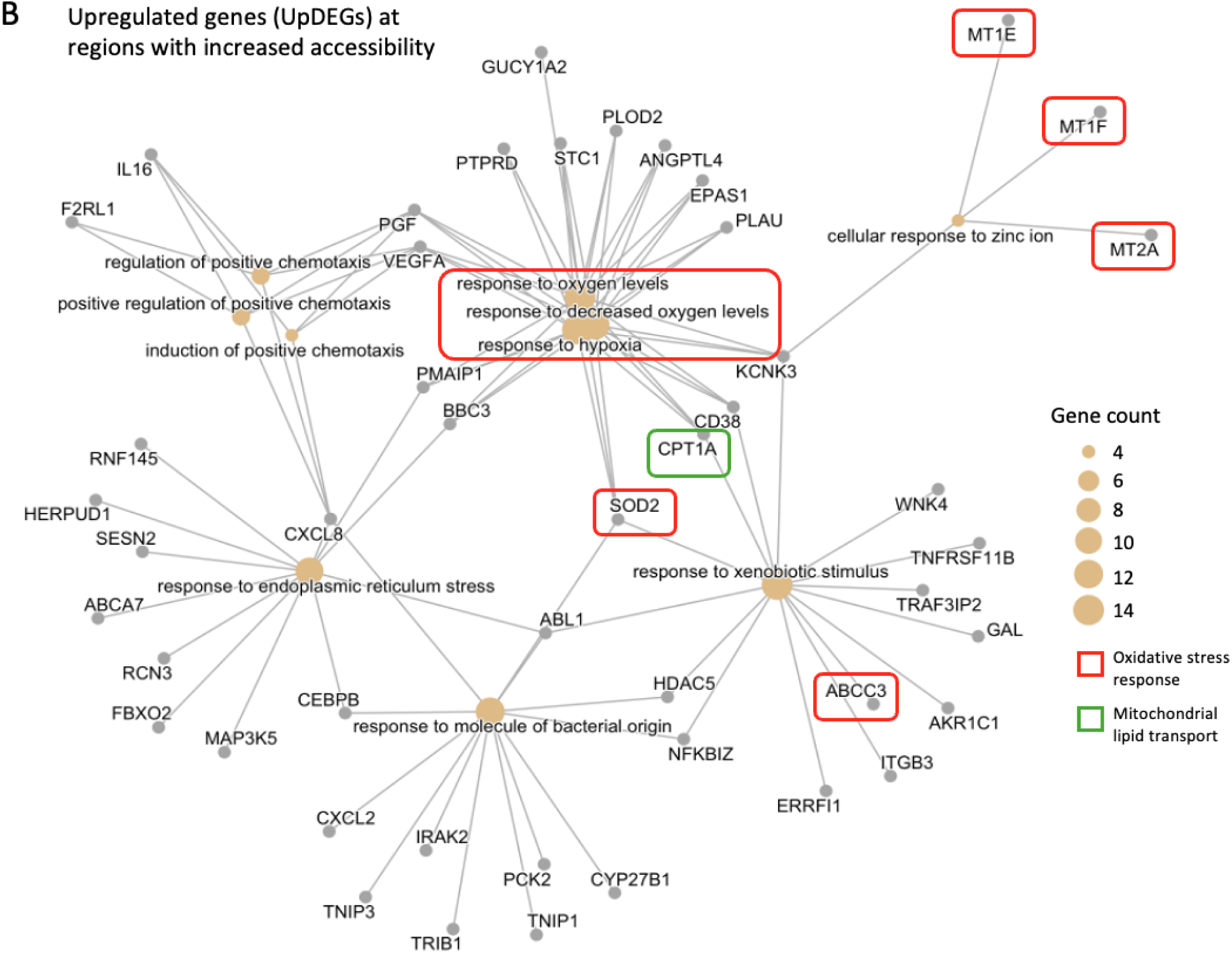
Chromatin accessibility remodeling and functional annotation of genes associated with the lipogenic state in TNBC cells. (A) Distribution of differentially accessible regions (DARs) across genomic features stratified by gene expression class. Schematic (top) indicates genomic regions relative to the transcription start site (TSS), including promoter (up to 3kb upstream), gene body, downstream, and distal regions. Bar plots show the percentage of genes within each class, upregulated (UpDEGs), downregulated (DownDEGs), or unchanged, that contain at least one DAR in the indicated region. DARs are separated into regions with increased accessibility (UpDARs, top) or decreased accessibility (DownDARs, bottom). (B) CNET plot showing genes associated with the top enriched biological pathways among upregulated genes (UpDEGs) that colocalize with regions of increased chromatin accessibility (UpDARs). Only genes linked to the displayed enriched Gene Ontology categories are included.

Functional enrichment analysis of UpDEGs associated with UpDARs revealed overrepresentation of stress-adaptive pathways, including responses to hypoxia, oxidative stress, xenobiotic stimuli, and endoplasmic reticulum (ER) stress (**Fig. 5B**). Network-based visualization (CNET plot) highlights the convergence of these pathways on key regulatory genes, including hypoxia-associated factors (e.g., *EPAS1*), chemotactic mediators (*VEGFA*, *IL6*), and oxidative stress response genes such as metallothioneins (*MT1F*, *MT1E*, *MT2A*) and *SOD2*. Genes involved in metabolic adaptation are also represented, including *CPT1A*, which mediates mitochondrial fatty acid import and supports fatty acid oxidation, and *ABCC3*, a transporter associated with xenobiotic export and cellular detoxification. These genes are connected to multiple enriched pathways, indicating coordinated activation of transcriptional programs that support redox balance, metabolic flexibility, and environmental adaptation. Notably, enrichment of metal ion homeostasis pathways, including cellular responses to zinc, further supports activation of mechanisms that mitigate oxidative stress. Together, these findings indicate that chromatin opening preferentially occurs at genes involved in adaptive cellular responses, consistent with selective enhancer activation rather than global epigenetic repression, while transcriptional repression is associated with comparatively limited and localized decreases in chromatin accessibility.

### The Adipocyte Secretome Supports Oxidative Capacity While Limiting ROS Accumulation in TNBC Cells

To determine whether adipocyte-induced stress-adaptive states extend to functional metabolism, we measured ROS levels and mitochondrial respiratory behavior. BT-549 cells were cultured in control unconditioned medium (UCM) or adipocyte-conditioned medium (ACM) for 72 hours, and oxidative stress was measured under basal conditions. Furthermore, to test whether reduced oxidative stress in ACM-treated cells depends on ongoing fatty-acid oxidation, we inhibited this process in a sample of ACM-cultured cells via treatment with etomoxir, an inhibitor of carnitine palmitoyltransferase-1 (CPT-1), to block mitochondrial uptake of long-chain fatty acids. Etomoxir was administered 48 hours prior to ROS measurements and maintained throughout the remainder of the experiment.

Following treatment, cells were harvested and stained with 2′,7′-dichlorodihydrofluorescein di-acetate (DCFDA) to quantify total intracellular reactive oxygen species, including hydroxyl and peroxyl radicals and peroxynitrite. Flow cytometric analysis using DCFDA revealed significantly lower global ROS levels in ACM-cultured BT-549 cells compared to UCM controls (**Fig. 6A**; *p* < 0.05). Inhibition of fatty-acid oxidation with etomoxir did not reverse this reduction, and ACM-treated cells continued to exhibit low ROS levels relative to UCM (*p* < 0.01). These results suggest that reduced oxidative stress in response to adipocyte exposure is not driven solely by ongoing fatty-acid oxidation.

**Figure 6.**
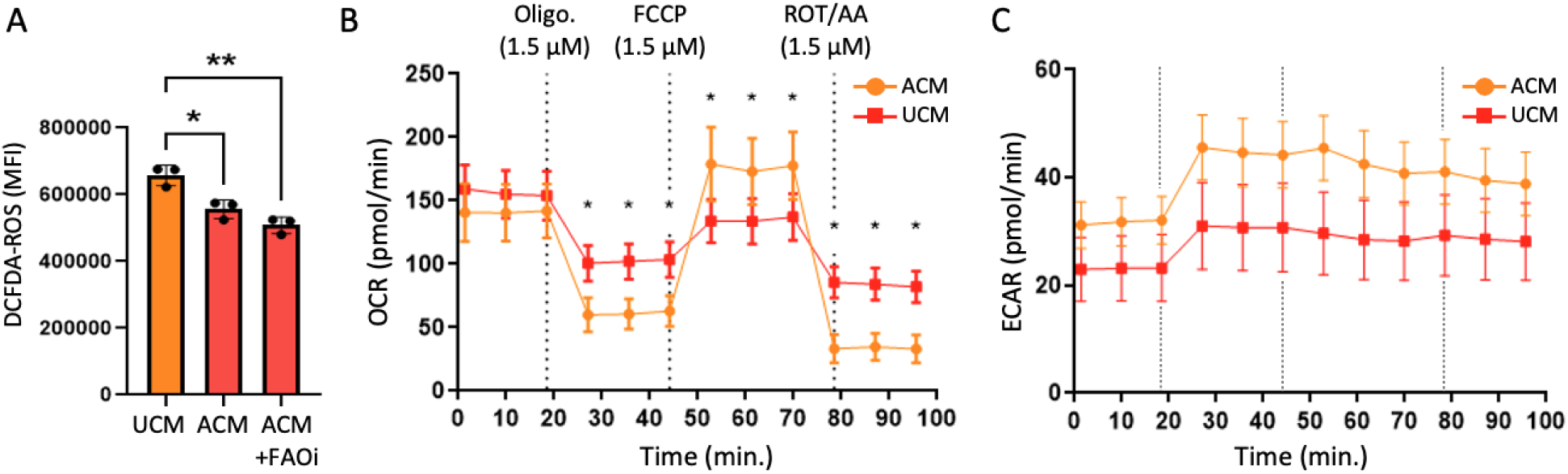
ACM-treated TNBC cells exhibit reduced oxidative stress and stress-adaptive metabolic responses. (A) Flow cytometric quantification of global reactive oxygen species (ROS) using the DCFDA probe in BT-549 cells cultured in control medium (UCM), adipocyte-conditioned medium (ACM), or ACM with fatty-acid oxidation inhibition (ACM + FAOi). Mean fluorescence intensity (MFI) is shown (*n* = 3 biological replicates). Statistical significance was assessed by one-way ANOVA (\**p* ≤ 0.05, \*\**p* ≤ 0.01). (B) Oxygen consumption rate (OCR) profiles measured during Seahorse XF Mito Stress Tests in BT-549 cells cultured in UCM or ACM for 72 hours. Sequential injections of oligomycin (ATP synthase inhibitor), FCCP (uncoupler), and rotenone/antimycin A (complex I and III inhibitors) are indicated by dashed vertical lines. ACM-treated cells exhibit comparable basal OCR, greater oligomycin-induced suppression, and increased FCCP-stimulated maximal respiratory capacity relative to UCM controls (*n* = 12 wells per condition; \**p* ≤ 0.05). (C) Extracellular acidification rate (ECAR) recorded simultaneously during the Mito Stress Test shown in (B). ACM-treated cells display elevated ECAR throughout the assay, indicating increased proton production during mitochondrial perturbation. Data are shown as mean ± SEM.

To assess how adipocyte secreted signals influence mitochondrial respiratory behavior, we performed Seahorse XF Mito Stress Tests on BT-549 cells cultured in control medium (UCM) or ACM for 72 hours. Basal oxygen consumption rates (OCR) were comparable between UCM-and ACM-treated cells prior to perturbation (**Fig. 6B**), indicating that adipocyte exposure does not increase steady-state mitochondrial respiration. Upon inhibition of ATP synthase with oligomycin, OCR decreased in both conditions, with ACM-treated cells exhibiting a significantly greater reduction compared to UCM controls (**Fig. 6B**; *p* < 0.05). This response suggests altered ATP-linked respiratory contributions in ACM-treated cells. In contrast, uncoupling of the electron transport chain with FCCP revealed a significantly higher maximal OCR in ACM-treated cells relative to UCM (**Fig. 6B**; *p* < 0.05), consistent with increased spare respiratory capacity. Following inhibition of complexes I and III with rotenone/antimycin A, residual OCR was reduced in both UCM and ACM-treated cells. Extracellular acidification rate (ECAR), recorded simultaneously during the Mito Stress Test, was elevated in ACM-treated cells throughout the assay relative to UCM controls (**Fig. 6C**), indicating increased proton production during mitochondrial perturbation. Together, these data indicate that adipocyte signaling reprograms mitochondrial function to enhance bioenergetic flexibility, preserving maximal oxidative capacity while altering ATP-linked respiration and acidification dynamics, consistent with a stress-adaptive metabolic state (**Fig. 7**).

**Figure 7.**
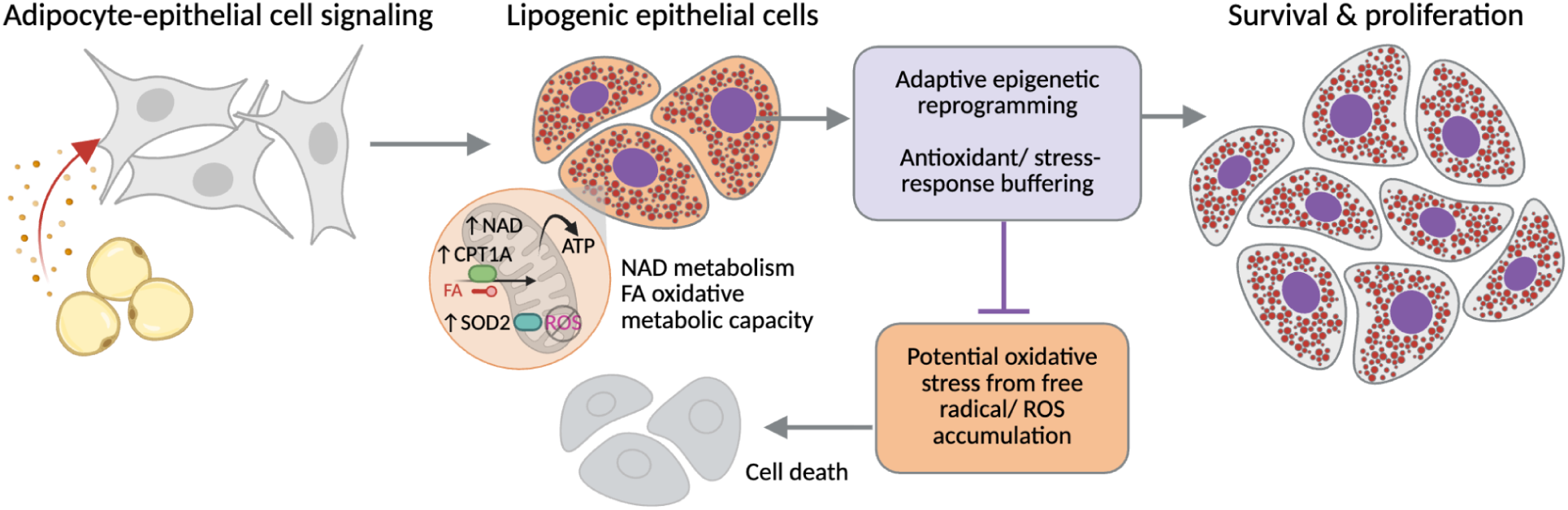
Adipocyte-induced lipogenic state promotes adaptive epigenetic reprogramming and oxidative stress buffering in TNBC cells. Adipocyte-derived signals induce a lipogenic state in TNBC epithelial cells accompanied by metabolic rewiring, including increased NAD metabolism and enhanced oxidative metabolic capacity. This state is associated with upregulation of pathways linked to fatty acid oxidation (e.g., CPT1A) and antioxidant defense (e.g., SOD2), consistent with transcriptomic and chromatin accessibility changes. These metabolic features are expected to generate reactive oxygen species (ROS) and impose oxidative stress. However, adaptive epigenetic reprogramming and antioxidant stress-response pathways limit ROS accumulation, enabling cell survival and proliferation. Cells that fail to sufficiently buffer oxidative stress undergo cell death.

## Discussion

In this study, we investigated how adipocyte-derived signals reshape metabolic–epigenetic coupling in triple-negative breast cancer (TNBC) cells by integrating transcriptomics, metabolic modeling, chromatin accessibility profiling, and functional metabolic assays. We find that adipocyte-conditioned media induces a durable proliferative phenotype following transient exposure, indicating that adipocyte signaling can establish persistent cellular states rather than transient responses. Systems-level modeling and RNA-seq analyses revealed metabolic rewiring characterized by enhanced NAD-linked metabolism and enrichment of redox-associated pathways, while predicted changes in canonical epigenetic cofactors were comparatively modest. These findings suggest that early lipogenic reprogramming is not driven by large shifts in bulk metabolite availability, but instead reflects redistribution of metabolic fluxes and adaptive responses to oxidative stress.

A key finding of this work is that adipocyte-induced metabolic rewiring is coupled to asymmetric chromatin remodeling, characterized by a strong bias toward increased chromatin accessibility. ATAC-seq analysis revealed that the majority of differentially accessible regions exhibit modest gains in accessibility, primarily within gene-proximal regions, and are reproducibly associated with transcriptional activation. Genes linked to these regions are enriched for stress-adaptive and antioxidant pathways, including hypoxia response, ER stress, and ROS buffering programs, such as metallothioneins and antioxidant enzymes such as SOD2 (48,49). Notably, increased accessibility was also observed at loci without immediate transcriptional changes, suggesting that chromatin opening may precede or poise future gene activation. Together, these findings indicate that lipogenic signaling preferentially promotes chromatin environments permissive for adaptive transcriptional programs rather than inducing widespread epigenetic repression.

In contrast, transcriptional repression in lipogenic TNBC cells appears to occur with minimal accompanying loss of chromatin accessibility. Downregulated genes showed limited association with decreased accessibility, and locus-specific analysis (e.g., *PARVA*) suggests that repression may occur within chromatin contexts that are already relatively constrained. These observations are consistent with a model in which gene silencing is mediated through regulatory mechanisms that do not require large-scale chromatin closure, such as transcription factor dynamics, cofactor competition, or local chromatin features. Importantly, these results do not exclude a role for epigenetic repression, but instead indicate that at early stages of lipogenic reprogramming, repression is not broadly encoded through global accessibility changes and may emerge through more targeted or temporally delayed mechanisms.

Functionally, adipocyte-exposed TNBC cells exhibit a bioenergetically flexible, stress-adaptive phenotype characterized by increased spare respiratory capacity, altered ATP-linked respiration, elevated extracellular acidification, and reduced ROS accumulation. These features are consistent with a metabolic state that balances increased oxidative capacity with mechanisms that buffer oxidative stress, a feature increasingly recognized as a hallmark of metabolic adaptation in cancer cells (50). Together with the chromatin and transcriptomic data, our findings support a model in which adipocyte-driven metabolic rewiring enables TNBC cells to tolerate the oxidative burden of lipid-rich environments by selectively activating stress-adaptive gene programs while maintaining regulatory capacity for gene repression. This adaptive coupling between metabolism and chromatin state reveals a mechanism by which the adipocyte-rich tumor microenvironment can promote survival and proliferation, and highlights metabolic-epigenetic vulnerabilities that may be leveraged for therapeutic intervention, including pathways linked to lipid oxidation and redox balance that have been implicated in therapy resistance (49,51), and metabolic enzymes such as ACSS2 that support chromatin regulation under stress conditions (52).

## Materials and Methods

### Cell culture

Media for each cell line is described in **Supplemental Methods**. BT-549 cells (ATCC #HTB-122) were grown at 37°C, with 5% CO_2_ in a humidified incubator. 1X DPBS without calcium and magnesium (Corning #20-031-CV) and 0.25% trypsin-EDTA (Thermo #25200056) was used for cell washing and harvesting. For ACM treatments, concentration was adjusted based on assay requirements, with 50% ACM used for transcriptomic and chromatin profiling experiments to preserve cell viability and enable detection of early regulatory changes, and 100% ACM used in clonogenic assays to assess maximal effects on long-term proliferative capacity. For long term storage, ∼1 - 2x10^6 cells were frozen in 1 mL 10% DMSO in FBS (Thermo #A4736401) at -80°C for 2 days, then transferred to -150°C (liquid nitrogen).

### Colony formation

BT-549 cells were treated with either control media or 100% ACM and incubated at 37°C for 72 hours before cells were harvested with 25% trypsin-EDTA (Thermo #25200056) and seeded in 6-well plates at the following cell densities: 150, 300, 600. The constant group denotes cells that were treated with the same media they were previously cultured in. The pre-treated condition denotes cells that were switched to control media after the 72 hour pre-treatment. Colonies were grown at 37°C for 3 weeks before cells were washed with 1X PBS (Corning #20-031-CV), fixed with methanol, stained with crystal violet 0.5% (wt/vol) in H2O, and imaged. Bar graphs were made using GraphPad Prism.

### RNA-seq

RNA-seq data from BT-549 TNBC cells cultured in unconditioned media (UCM) or 50% adipocyte-conditioned media (ACM) for 24 hours (*n* = 3 per condition) were obtained from our previously published study (29), and are available online (NCBI GEO GSE282461). In that study, RNA was extracted, and libraries were prepared and sequenced (paired-end, 150 bp; Q30 ≥ 85%) by Novogene. Sequence alignment, transcript quantification, and differential expression analysis were performed using STAR, RSEM, and DESeq2, respectively, as previously described. For the present study, processed gene expression outputs were used directly for downstream analyses and integrated with newly generated ATAC-seq data from matched sub-samples.

### Genome-scale metabolic modeling and flux balance analysis

Genome-scale metabolic models were constructed using the Recon3D framework, incorporating transcriptomic data and biochemical constraints to improve physiological relevance. Enzyme capacity constraints were estimated using kcat values from the BRENDA database (53) and inferred enzyme abundances based on RNA-seq data, following established approaches (54).

Thermodynamic directionality constraints were applied using standard transformed Gibbs free energies (ΔG′°) from the Virtual Metabolic Human (VMH) database (55,56). Flux balance analysis (FBA) was performed using the VMH biomass reaction as the objective function to estimate cellular growth potential. Models were simulated under adipocyte-conditioned media (ACM) and unconditioned media (UCM) conditions. The extracellular compartment was constrained to reflect the composition of DMEM/F-12 culture medium (Thermo Fisher Scientific #11320) supplemented with fetal bovine serum (FBS), ensuring consistency between *in silico* and *in vitro* experimental conditions.

### Statistical and pathway enrichment analysis

Reaction-level flux differences between ACM and UCM models (*n* = 3 per condition) were assessed using two-tailed Student’s t-tests with Benjamini-Hochberg correction for multiple testing (57). Significantly altered reactions (adjusted *p* < 0.05) were subjected to subsystem enrichment analysis using Fisher’s exact test, with the full Recon3D network as background (31).

### ATAC-seq

ATAC-seq was performed on matched sub-samples of the same BT-549 cell populations used for RNA-seq. Briefly, cells cultured in unconditioned media (UCM) or 50% adipocyte-conditioned media (ACM) for 24 hours (*n* = 3 per condition) were split at harvest, with parallel aliquots processed for RNA-seq (Fig. 2) and ATAC-seq (Fig. 3). Replicate IDs were maintained consistently across modalities (R1 - R3 for UCM and ACM) to preserve one-to-one correspondence between chromatin accessibility and transcriptional profiles. Frozen cell pellets were submitted to Novogene for ATAC-seq library preparation and paired-end sequencing. Raw reads were trimmed using Trim Galore and aligned to the human reference genome (GRCh38/hg38) using BWA-MEM (58). PCR duplicates were removed using Picard. To account for Tn5 transposase insertion bias, aligned reads were offset by +4 bp on the positive strand and -5 bp on the negative strand. Peaks were called using MACS3 (59), and differential accessibility analysis was performed using DiffBind with an FDR threshold of < 0.05. Peaks were annotated to genomic features using ChIPseeker. Differentially accessible regions (DARs) were linked to associated genes and intersected with RNA-seq results to identify genes exhibiting coordinated changes in chromatin accessibility and expression. Functional enrichment analysis and gene-concept network (CNET) visualization were performed in R using clusterProfiler (60), with intermediate data organization and filtering steps performed in Microsoft Excel (v.16).

### Western blot

BT-549 cells were harvested after 72 hours of ACM treatment and lysed with RIPA Buffer with 5% Phosphatase inhibitor to harvest protein content. The cytoplasmic and nuclear protein fractions were isolated using the Pierce™ NE-PER® Nuclear and Cytoplasmic Extraction Reagent Kit (Thermo Scientific #78833). 15-20 g of protein with 1X MOPS SDS (Invitrogen #NP0001) and 5 μL of Precision Plus Protein Kaleidoscope ladder (BioRad #1610375) were run in a 12% Bis Tris gel (Invitrogen #NP0321BOX), transferred onto a nitrocellulose transfer blot (BioRad #1704158), and blocked with 3-5% BSA in TBST (pACSS2, ACSS2, H3K9ac, H3K27ac) or 3-5% Milk in TBST (B-Actin). The antibodies used were as follows: ACSS2(Phospho-Ser659) (Signalway #58003), ACSS2 polyclonal (Thermo Fisher #PA5-26612), Acetyl-histone H3 (lys9) (Cell signaling Technology #9649T), and Anti-histone H3 (acetyl K27) (Abcam ab4729). Secondary antibodies with HRP conjugation and Supersignal West Femto Maximum Sensitivity Chemiluminescent Substrate (Thermo Fisher #34094) were used to visualize bands with the Agilent iBright imager. Bands were analyzed using ImageJ where the background signal (mean intensity) was subtracted from the target band (mean intensity) and the control band (mean intensity), then the normalized target band value was divided by the normalized control band value.

### Immunostaining and confocal imaging

BT-549 cells were seeded onto poly-L-lysine-coated coverslips (Millipore Sigma #P4707) and cultured overnight prior to treatment with unconditioned media (UCM) or adipocyte-conditioned media (ACM) for 48 hours. Cells were fixed with 4% paraformaldehyde (Thermo Fisher Scientific #J61899-AK), permeabilized with 0.1% Tween-20 (VWR #76347-662), and blocked with 5% BSA (Cell Signaling Technology #9998S). Primary antibodies ACSS2(Phospho-Ser659) (Signalway #58003) were applied at 1:500 dilution, followed by Alexa Fluor 488-conjugated goat anti-rabbit IgG secondary antibody (Abcam, #ab150077). Nuclei were counterstained using SiR-DNA (Cytoskeleton, #CY-SC007). Coverslips were mounted in glycerol-based medium and imaged using a Leica Stellaris SP8 confocal microscope at 40× magnification. Additional experimental details are provided in the Supplementary Methods.

### Flow cytometry

BT-549 TNBC cells were seeded at a density of 1 × 10^5 cells per well in a 6-well plate containing 2 mL of complete medium and grown overnight. Drug-treated groups were treated with etomoxir for 48 hours, then all cells were cultured in either UCM or ACM for 72 hours. Drug-treated groups were given additional dosages of etomoxir for the remainder of the experiment. After 72 hours, cells were pelleted and stained with 100 μM of DCFDA/H2DCFDA (Abcam ab113851) or 150 nM of MitoTrackerRed (Thermo #M22425) for 30 minutes at room temperature. Buffer solutions and the positive control TBHP drug was prepared as directed. Cells were washed three times with FACS Buffer and analyzed using filters compatible with 485/535 nm (APC) and 644/665 nm (FITC), respectively, on a CytoFLEX Flow Cytometer with CytExpert software. Data analysis was completed using FlowJo software and bar graphs were generated using GraphPad Prism.

### Seahorse XFe24 flux analysis

BT-549 cells were cultured in ACM or UCM for 72 hours prior to seeding ∼5x10^3 cells per well in compatible microplates for the Seahorse XFe24 Flux Analyzer (Agilent, Santa Clara, CA). Additional 96-well plates were seeded in parallel to measure viable cell with Cell Titer Glo luminescence (VWR #G7571) on a BioTek plate reader. The Mito Stress Test (Agilent #103015-100) protocol was conducted as directed with the use of 1.5 µM of oligomycin, 0.5 µM of carbonyl cyanide-4 (trifluoromethoxy) phenylhydrazone (FCCP), and 0.5 µM of rotenone/antimycin A (AA) for OCR time course measurements.

## Author contributions

A.T. performed ACM treatments, cell culture, imaging, immunostaining (western blotting and immunofluorescence), flow cytometry, Seahorse assays, data analysis, and manuscript preparation. M.J. performed RNA-seq and ATAC-seq data processing and downstream analyses in R, with assistance from Y.W. S.H. and M.K. performed metabolic modeling, flux balance analysis, and pathway enrichment analysis. A.I., M.D., and C.J.H. contributed to identification of key mediators and interpretation of metabolic function. M.K., C.J.H., and K.A.H. conceived and designed the study. K.A.H. performed additional data analyses, edited figures, and finalized the manuscript.

## Acknowledgements

This study was supported by Developmental Funds from the Winship Cancer Institute of Emory University. We thank the following Emory collaborators for generously providing reagents, assays, and specialized support: M. Lee (ACM treatments); M. Shamugam (Etomoxir); S. Kang, A. Marcus, and colleagues (Seahorse FX Analyzer materials); Y. Wan (colony formation experiments). The research was also supported in part by the Pediatrics/Winship Flow Cytometry Core of Winship Cancer Institute of Emory University, Children’s Healthcare of Atlanta and NIH/NCI (award P30CA138292), and the Emory University Integrated Cellular Imaging Core Facility (RRID:SCR_023534). The content is solely the responsibility of the authors and does not necessarily represent the official views of the National Institutes of Health.

