## Supplemental Figures and Methods for "Early Epigenetic and Metabolic Responses to the Adipocyte Secretome Reveal Stress-Adaptive States in Triple-Negative Breast Cancer"

- **Figure S1.** Chromatin accessibility at the transcriptionally repressive *PARVA* locus.
- **Figure S2.** Single cell immunofluorescence cytometry profiles of pACSS2 in BT-549 cells.

### Supplemental Materials and Methods

- Preparation of cell culture media
- Immunostaining and confocal imaging (detailed methods)

### SUPPLEMENTAL FIGURES

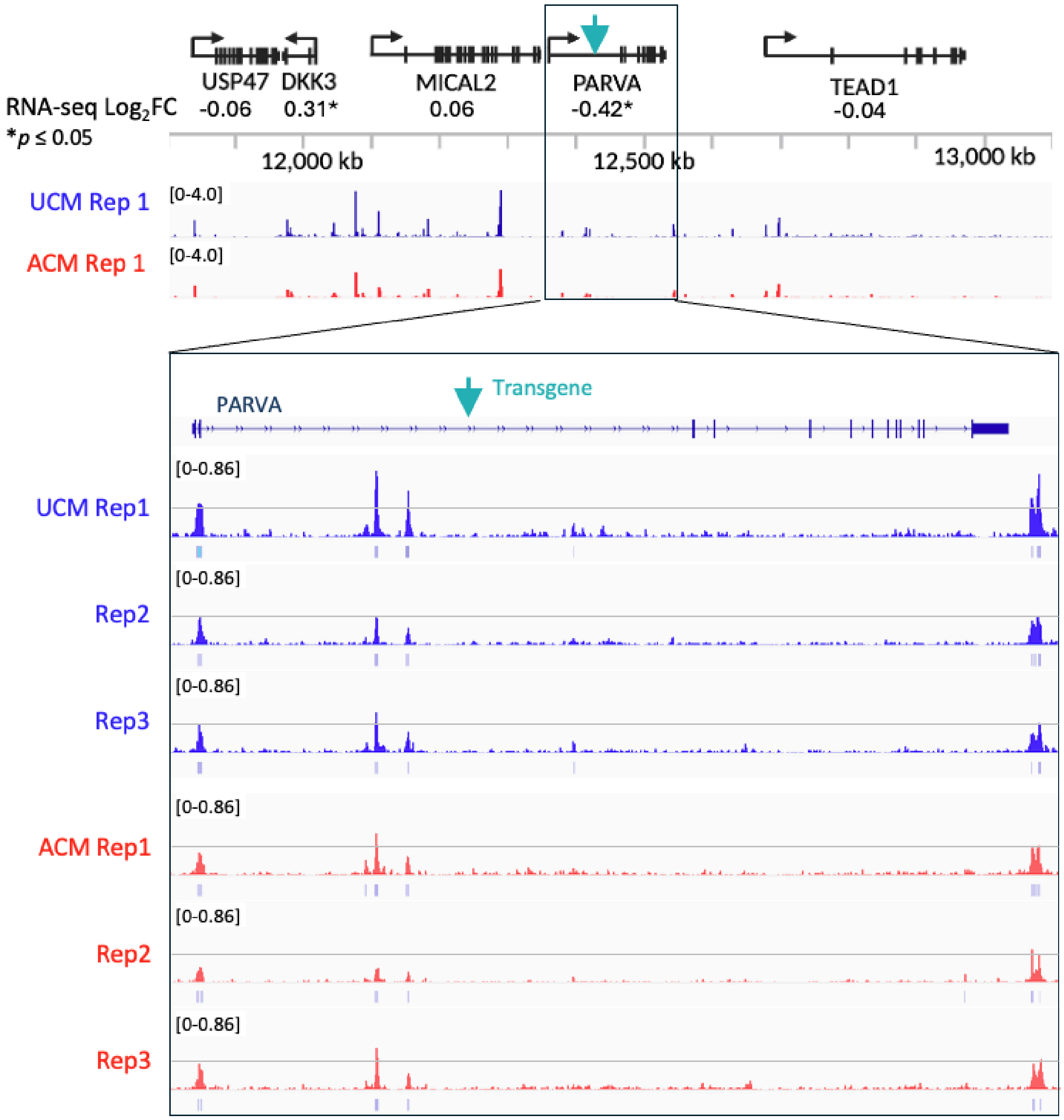


**Figure S1. Chromatin accessibility at the transcriptionally repressive PARVA locus.** Genome browser view of the *PARVA* locus and neighboring genes. The upper panel shows gene models and RNA-seq $log_2$ fold change values. Middle and lower panels show representative ATAC-seq signal tracks for UCM (blue) and ACM (red) conditions. The transgene insertion site is indicated. The *PARVA* locus exhibits low baseline accessibility with modest additional reduction upon ACM treatment.

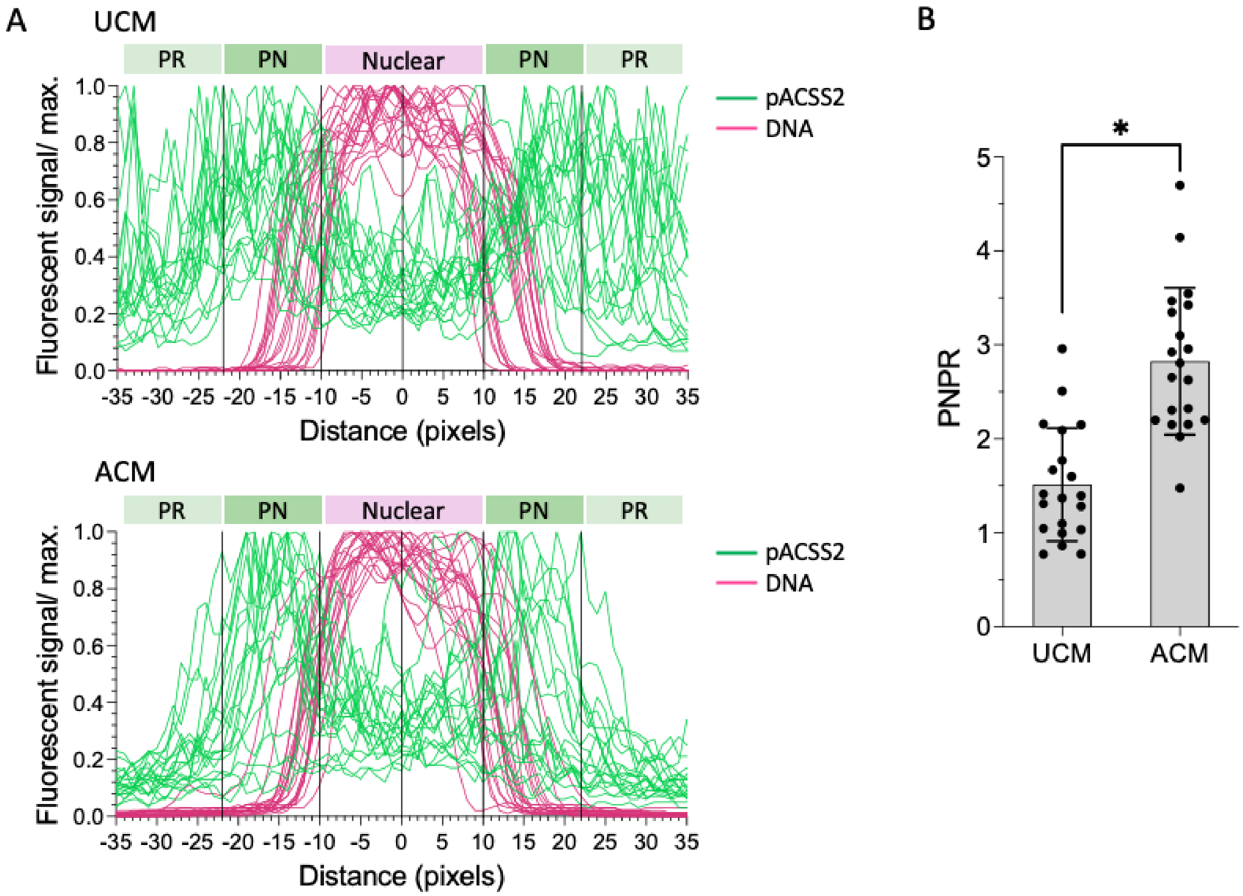


**Figure S2.** Single cell immunofluorescence cytometry profiles of pACSS2 in BT-549 cells. (A) Fluorescence profiles for 20 cells per condition (UCM or ACM). Overlay, pACSS2, and DNA channels were imported into FIJI/ImageJ2 (version 2.14.0/1.5f). In the overlay image, the selection tool was used to draw a freehand line (70px, 51.1 μm long, 10px thickness) centered at the nucleus. The selection was restored in the pACSS2 and DNA channels, and the Plot Profile analysis tool was used to measure intensity at each point. (B) Perinuclear to peripheral ratios (PNPR) represent the distribution of the pACSS2 signal outside of the nuclear zone (DNA profile peak, -10 to 10px). Perinuclear (PN) = mean(pACSS2 at -22 to -11px and 11 - 22px). Peripheral (PR) = mean(pACSS2 at -35 to -23px and 23 - 35px). PNPR = PN / (PR + 0.0001). *$p < 0.001$

### SUPPLEMENTAL MATERIALS AND METHODS

#### Preparation of cell culture media

| Cell line (catalog no.) | Base medium | Formula |
|---|---|---|
| BT-549 (ATCC #HTB-122) | RPMI-1640 | 10% FBS, 0.8 µg/mL insulin |
| HCC1806 (ATCC #CRL-2335) | RPMI-1640 | 10% FBS, 1% penicillin-streptomycin, 1% L-glutamine, 1% Sodium Pyruvate |
| 4T1 (ATCC # CRL-2539) | DMEM | 1% L-glutamine + 10% FBS + 1% sodium pyruvate + 1% non-essential amino acids + 0.01 mg/mL human recombinant insulin |

#### Immunostaining and confocal imaging (detailed methods)

Coverslips were sterilized with 70-100% ethanol for 10 minutes at room temperature, then one coverslip was placed in each well of a 6-well plate for further sterilization with ethanol. After washing twice with 1X PBS (Corning #20-031-CV), coverslips were coated with 500 µL of Poly-L-Lysine (Millipore Sigma #P4707-50ML) for 30 minutes to 1 hour at room temperature, followed by three 1X PBS rinses. Cells were seeded at 10^5 cells in 3 mL of growth medium on the Poly-L-Lysine coated coverslips before overnight incubation at 37°C. The next day, the medium was replaced with control UCM or adipocyte-conditioned media (ACM) for 48 hours. Cells were then washed with 1X PBS and fixed with 4% paraformaldehyde in PBS (Thermo Fisher Scientific #J61899-AK) for 10 minutes at room temperature. Permeabilization was performed with 0.1% Tween-20 in PBS (made using VWR #76347-662) for 5 minutes, followed by three washes with 1X PBS. After blocking the cells with 5% BSA in PBS (made using Cell Signaling Technology #9998S) for 1 hour, coverslips were incubated with primary antibody diluted 1:500 in 5% BSA for over 2 hours at 4°C. Secondary Goat Anti Rabbit IgG H&L (Alexa Fluor® 488) (ab150077) was centrifuged and diluted 1:500 in 5% BSA blocking buffer, then incubated for 40-50 minutes at room temperature in the dark. Following three 1X PBS washes, nuclei were counterstained with 2 µM SiR-DNA (Cytoskeleton #CY-SC007) in culture medium for 1 hour at 37°C and washed once with 1X PBS. Coverslips were mounted on glass slides using 50% glycerol/1X PBS (made using Sigma-Aldrich #G5516-100ML) and sealed with clear nail polish for imaging with a Stellaris SP8 confocal imager at 40x magnification.
